# Functional evolution of SARS-COV-2 Spike protein: adaptation on translation and infection via surface charge of spike protein

**DOI:** 10.1101/2022.05.16.492062

**Authors:** Xiaolong Lu, Gong Zhang

## Abstract

The SARS-COV-2 virus, which causes the COVID-19, is rapidly accumulating mutations to adapt to the hosts. We collected SARS-COV-2 sequence data from the end of 2019 to April 2022 to analyze for their evolutionary features during the pandemic. We found that most of the SARS-COV-2 genes are undergoing negative purifying selection, while the spike protein gene (S-gene) is undergoing rapid positive selection. From the original strain to the alpha, delta and omicron variant types, the Ka/Ks of the S-gene increases, while the Ka/Ks within one variant type decreases over time. During the evolution, the codon usage did not evolve towards optimal translation and protein expression. In contrast, only S-gene mutations showed a remarkable trend on accumulating more positive charges. This facilitates the infection via binding human ACE2 for cell entry and binding furin for cleavage. Such a functional evolution emphasizes the survival strategy of SARS-COV-2, and indicated new druggable target to contain the viral infection. The nearly fully positively-charged interaction surfaces indicated that the infectivity of SARS-COV-2 virus may approach a limit.

## 1. Introduction

The SARS-COV-2 virus has been causing COVID-19 pandemic for more than two years. During the pandemic, the virus is rapidly accumulating mutations. The three major variants, namely alpha, delta and omicron, caused significant waves of infection worldwide in early 2021, mid 2021 and early 2022. Each infection wave is much more serious than the initial outbreak in Wuhan, China around early 2020, and the infections in each wave increased dramatically [1]. With the mitigation of quarantine policies in many countries, the transmission will be booming, and the transmission between human and animals may become more frequent. Thus, omicron should not be the last variant of SARS-COV-2. The main evolutionary force of the viral mutations includes infectivity, reproduction efficiency and immune evasion. The earliest infectivity-increasing mutations were observed in China (V367F, W436R and D364Y), where strict isolation policies were deployed; in contrast, the other early mutations in the rest of the world did not show increased affinity of S-RBD and human ACE2 because of relaxed quarantine measures [2,3]. The worldwide massive vaccination was also a driving force of virus evolution. The major variants emerged soon after such massive vaccination, causing immune evasion and significantly reduces the effectiveness of vaccines [4,5]. However, little was known about the mutations on the reproduction efficiency, i.e. the adaptation on the viral protein synthesis.

The epidemiology indicated that the SARS-COV-2 virus is adapting human for co-existence due to the increasing infectivity and decreasing mortality. However, many studies showed that the SARS-COV-2 is under strong purifying selection [6,7]. This normally means that the functional mutations are being excluded, which is counteracting the adaptation [8]. Such a contradiction needs more clarification.

In this study, we focused on the translational adaptation of the SARS-COV-2 virus and the surface properties of the spike protein to provide more insights on the abovementioned questions, and depict the major trend of the SARS-COV-2 evolution during the evolution.

## 2. MATERIALS AND METHODS

### 2.1. Sequencing data analysis

The original RNA-seq and Ribo-seq data (Calu3 cells infected with SARS-COV-2 for 7h) were downloaded form Gene Expression Omnibus database (GEO) under accession number GSE149973[9], In brief, all raw sequencing data low quality reads and linker were removed using fastp[10]. For SARS-COV-2 gene expression quantification, cleaning reads were aligned to human refmRNA (UCSC) in order to remove human reads using Fanse3 program[11,12], after human reads removal, the remaining reads mapped to the Wuhan Hu-1 (NCBI Accession NC_045512.2. Mapping parameter, Ribo-seq: -S10 -E1 -U1, RNA-seq: -S12 -E2), reads per kilobase per Million mapped reads (RPKM) of each genes were calculated in golang program.

### 2.2. Proteomic and RNC-seq expression data

Proteomic (quantitative technique: DIA, A549 cells infected with SARS-COV-2 for 24h) and RNC-seq (human HBE cell line) expression data were extracted from two independent studies supplementary materials, respectively[13,14].

### 2.3. SARS-COV-2 genomes data analysis

SARS-COV-2 genomes for temporal evolution analysis were download form NCBI SARS-COV2 database at 2022/4/24 (https://www.ncbi.nlm.nih.gov/sars-cov-2/), with a list of 932520 SARS-COV-2 genomes, only sequences with length longer than 29500 and have zero N bases will be included in the subsequent analysis. Genomes for CAI and ITE evolution analysis were downloaded form GISAID database (www.epicov.org) at 2021/12/01.

Protein-coding sequences were exract by using Exonerate2.2 with affine:local model, (https://www.ebi.ac.uk/about/vertebrate-genomics/software/exonerate), which perform gapped alignment against a database, this model equivalent to the classic Smith-Waterman-Gotoh type of alignment. Lineage assignment using Pangolin lineage command line tool v3.0 (https://github.com/cov-lineages/pangolin) with standard model[15].

### 2.4. Ka/Ks and charge evolution analysis

Due to high closely related (identity ∼ 99%) of SARS-COV-2 genomes, we used MAFFT v7 as multiple sequence aligner[16]. Each cds Ka/Ks calculation was performed by Ka/Ks calculator2.0 using YN method[17,18]. For charge evolution, Protein translation and net charge calculations were performed using gotranseq v0.3.0 (https://github.com/feliixx/gotranseq) and peptides v1.2.2 (https://github.com/dosorio/Peptides)[19], respectively.

### 2.5. Codon adaptation analysis

CAI (Codon adaptation index), ITE (Index of Translation Elongation) and RSCU (Relative Synonymous Codon Usage) were calculated using DAMBE7, CAI is a measure of the host codon adaptation for a gene sequence. ITE it is similar to CAI, which incorporates both tRNA-mediated selection and background mutation bias and fits protein production better than CAI[20–22].

### 2.6. Electrostatic energy and surface electrostatic potential calculations

The complex structure of the SARS-COV-2 RBD and ACE2 complex (PDB: 6M0J), SARS-COV-2 Spike with furin cleavage site structure (PDB: 7FG7) and Human furin structure (PDB: 5MIM) were obtained from PDB database.

The three-dimensional structures of three variants (alpha, delta and omicron) were generated by in silico mutation using UCSF chimera software v1.15 (Dunbrack backbone-dependent rotamer library method)[23,24], all site mutations selected the most probable isoforms. PyMol was used to calculate the surface electrostatic potential by Adaptive Poisson Boltzmann solver (APBS), the color scale range was set from −1.0 to 1.0 kT/Å.

We used delphi force webtool (http://compbio.clemson.edu/delphi-force/) to calculate the electrostatic energy of the RBD and ACE2 binding domain, the required PQR file was generated by delphiPKa (http://compbio.clemson.edu/pka_webserver/)[25,26]. DelphiPKa calculation was used charmm force field, pH and salt concentration were set to 7 and 0.15M, respectively, other setting uses standard mode, delphiforce calculation uses the Gaussian-smoothed mode, the Grid Resolution uses 2.0Å, and the salt concentration is set to 0.15M.

## 3. Results

### 3.1. The S-gene is the only gene undergoing stepwise positive selection

We used Ka/Ks to evaluate the selection of the SARS-COV-2 sequences from the end of 2019 to April 2022. Ka/Ks >1 means that the gene is undergoing positive selection and drives the adaptation. Ka/Ks <1 indicates purifying selection, which stabilizes the gene. It is noted that the Ka/Ks of the S-gene was below 1 till February 2021, showing that in the first year of pandemic, S-protein was under purifying selection in general, indicating that the selective pressure was not that significant. However, the Ka/Ks surpassed 1 from Feburary 2021 and is constantly increasing, showing that the S-protein is undergoing an increasingly stronger positive selection (Fig. 1A).In contrast, the other important and highly expressed genes, such as the N-gene and M-gene as structural genes, the orf1ab gene for viral replication, the orf3a and orf7a genes that mediates cell inflation, were undergoing purifying selection, because their Ka/Ks were almost constantly below 1. The Ka/Ks of N-gene was transiently above 1 between February and September 2021 and then dropped back below 1, suggesting that the mutations generated in this period were not essential to maintain pandemic. These results showed that the S-gene was the only SARS-COV-2 gene that underwent positive selection. Such positive selection may be driven by the massive vaccination worldwide, because most COVID-19 vaccines include or produce S-protein.

**Figure 1.**
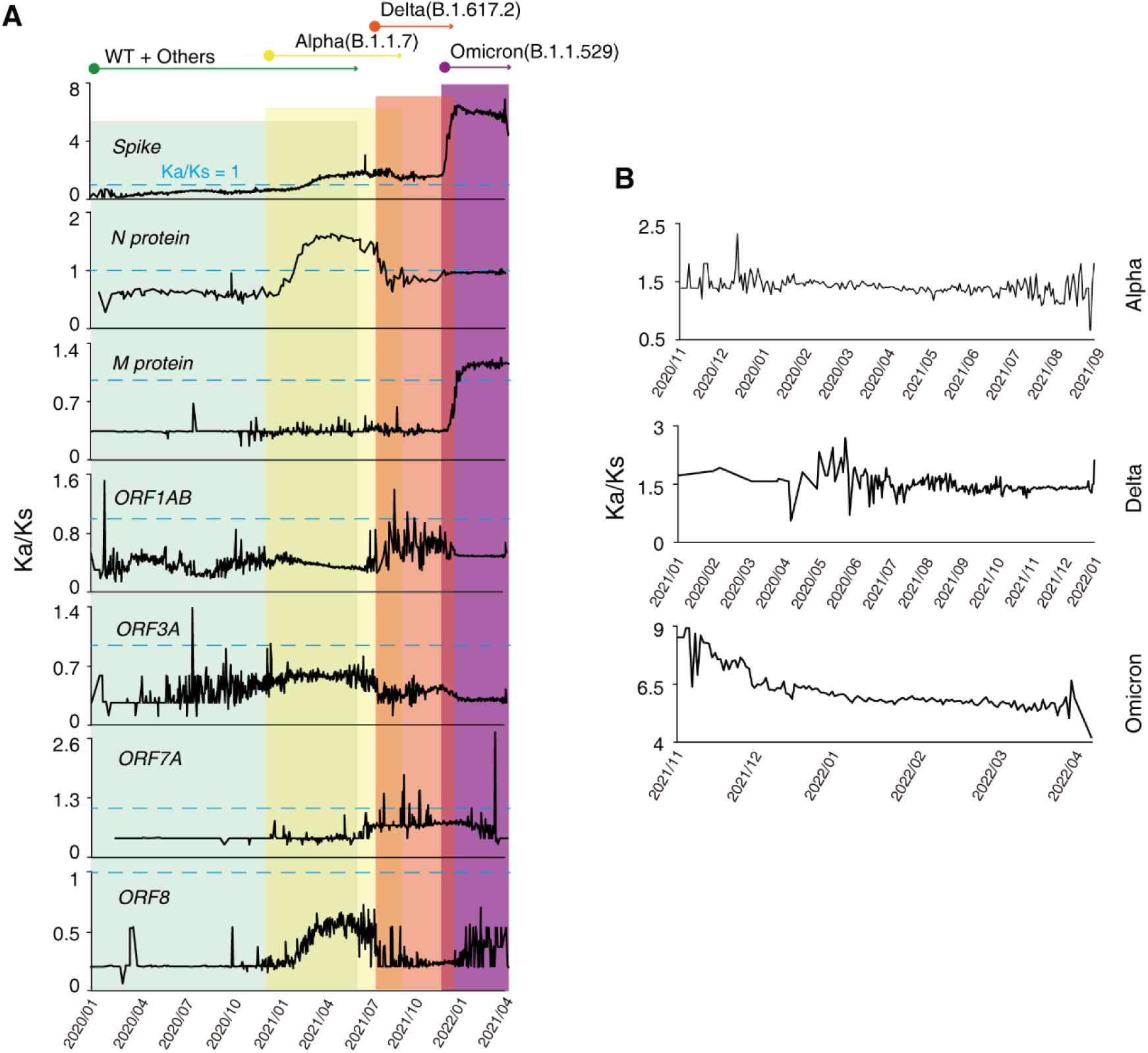
Ka/Ks analysis of SARS-COV-2 evolution. Ka/Ks of each gene was calculated using the SARS-CoV-2 genome sequences. The data was averaged in a day-wise manner. (A) Temporal Ka/Ks analysis of annotated canonical CDS. The background colors indicate the major lineages (Wild-type, alpha, delta and omicron) and their approximate pandemic timespan. E protein, ORF7B, ORF6 and ORF10 were not shown due to insufficient data. Cyan dashed line showed the threshold of Ka/Ks=1. (B) Ka/Ks evolutionary trends of major variants.

Interestingly, the stepwise elevation of the positive selection coincides temporally with the major SARS-COV-2 variation types (alpha, delta, omicron. Fig. 1A). However, within one variant type, the Ka/Ks of S-gene slightly decreased (Fig. 1B). This indicated that the mutations were accumulated stepwise.

### 3.2. SARS-COV-2 codon usage did not evolve towards more efficient translation

Adaptation to the host translation system is needed for efficient viral protein production and replication of the virus. Such adaptation is largely represented by similar codon usage in the host cells. Using the CAI (codon adaptation index) and ITE (index of translation elongation) to represent the codon adaptation, we showed that the SARS-COV-2 genes with higher CAI and ITE were expressed in a significantly higher level (Fig. 2A). This trend was also visible in Ribo-seq data (Fig. 2B), but CAI and ITE do not correlate to mRNA expression level at all (Fig. 2C). These results showed that the protein expression preference towards the codon adaptation happened indeed in the translation process of SARS-COV-2. However, SARS-COV-2 CAI and ITE were significantly lower than that of the human cells and other well-known human viruses, respectively, while the human viruses have comparable CAI and ITE against the human cells (Fig.2D). Here, we used the genomic data to calculate the CAI and ITE for all virus, and we used the RNC-seq (sequencing of the translating mRNAs) data of the human lung cell line A549 to reflect the actual codon demand in human lung cells, which was the primary infection host of SARS-COV-2. These results showed that the SARS-COV-2 were not fully adapted to human in terms of codon usage.

**Figure 2.**
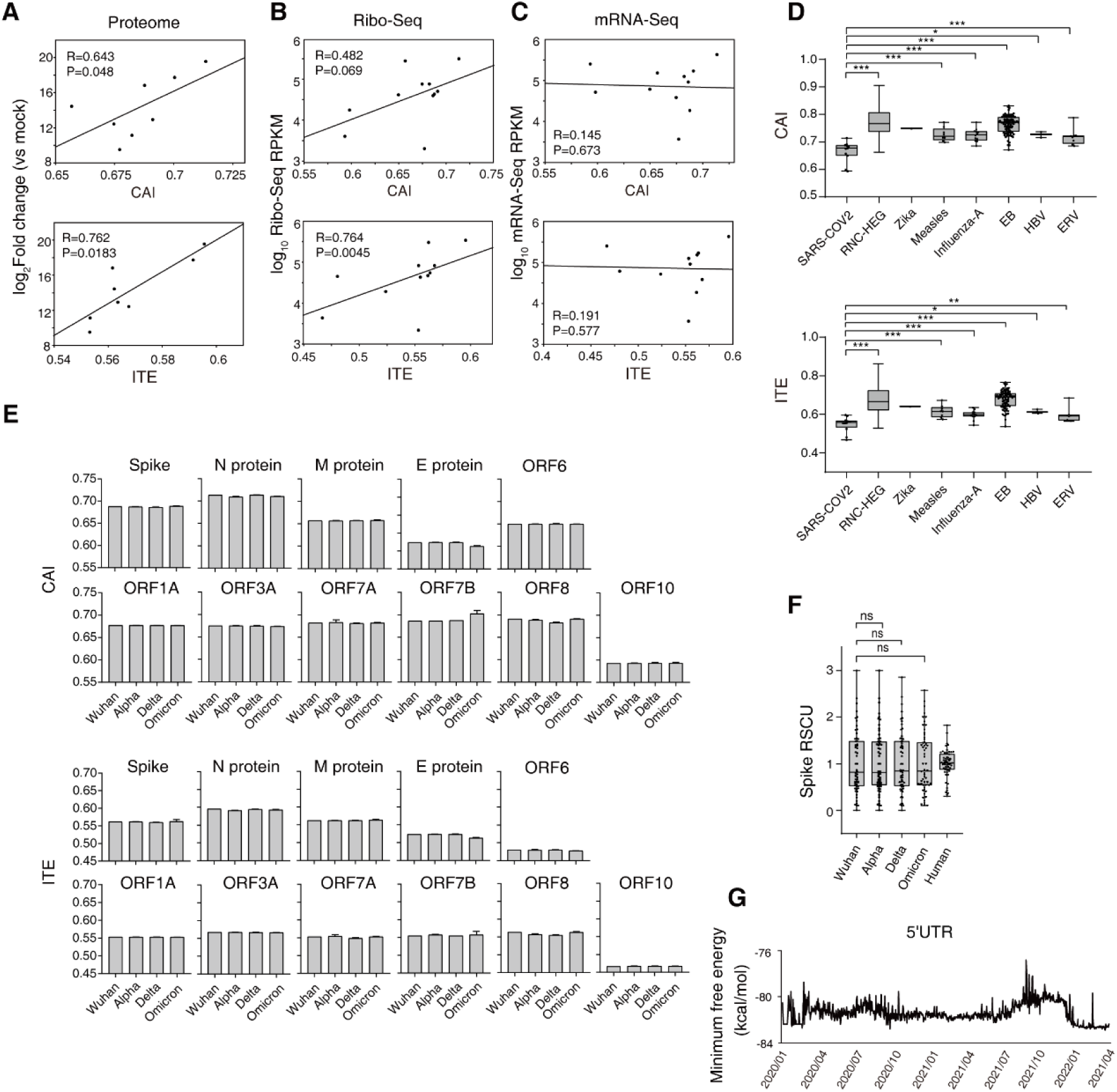
SARS-COV-2 codon adaptation analysis. (A, B, C) Correlation of CAI/ITE values against the proteome proteomic, Ribo-seq and RNA-seq gene expression level in SARS-COV-2 infected cells. Spearman correlation coefficient. (D) CAI and ITE value calculated for SARS-COV-2 genes, RNC-HEG (translating highly expressed genes), common human viruses and ERV (human endogenous retroviruses) sequence. (E) CAI and ITE values of each gene grouped by variant. (F) Box plot of Spike protein RSCU values grouped by variant. Human RSCU was compared as negative control. (G) Evolutionary trends of SARS-COV2 5’UTR MFE.

To assess the evolutional trend of the codon adaptation of SARS-COV-2 during the pandemic, we calculated the CAI and ITE of the typical strains of the wild-type (the original strain in Wuhan) and the major variants alpha, delta and omicron (Fig. 2E). Most genes showed a constant CAI and ITE during the pandemic. Only the orf7b gene showed an increase in CAI and ITE. However, the orf7b was lowly expressed, indicating the limited biological impact of its increase in codon adaptation.

The S-gene, which is the only gene underwent positive selection, showed a decrease in CAI and ITE from the original strain to delta variant, and minimally increased in omicron variant. Such a minimal increase may not cause any visible biological effect. We also calculated the RSCU of the S-gene of all collected virus sequences. Compared to condensed RSCU distribution of human genes, S-genes of all major variants showed a wide scatter of S-gene RSCU values (Fig. 2F), indicating that the optimal codon usage was not a selective force during the pandemic.

Translation efficiency is determined by translation initiation and translation elongation, while CAI, ITE and RSCU can only reflect translation elongation efficiency. According to previous research, 5’-UTR minimum free energy (MFE) characterizes translation initiation efficiency. We calculated the SARS-COV-2 5’UTR MFE between end of 2019 to April 2022, it can be seen that there was no continuous upward trend in two years 5’utr evolution (Fig. 2G), and there was only a brief and small increase before July 2021 and January 2022, indicating that the translation initiation efficiency did not have sign of enhancement.

In sum, the SARS-COV-2 virus did not mutate towards optimal translation efficiency. We need to investigate the evolutional advantage from other aspects.

### 3.3. S-gene accumulated positive charges during pandemic

Electrostatic force is a major force to guide protein-protein interactions. Compared to van der Waals force, the electrostatic force decays much slower against distance, which facilitates the binding of virus to other proteins in various important processes. We analyzed the changes of charges (at pH=7) and found similar pattern as the Ka/Ks analysis. The S-gene showed a steady increase in positive charge, and this increase was remarkably speeded up since June 2021, which was the time when delta variant was leading the pandemic. The charges of other genes did not apparently change during the pandemic in general (Fig. 3A). The accumulation of the positively charged amino acid residues were mainly found in the RBD and furin domains (Fig. 3B). The net charge in RBD domain increased from 1.5 to 4, and the net charge in furin domain increased from 3 to 4. These results indicated that these changes may be functionally meaningful.

**Figure 3.**
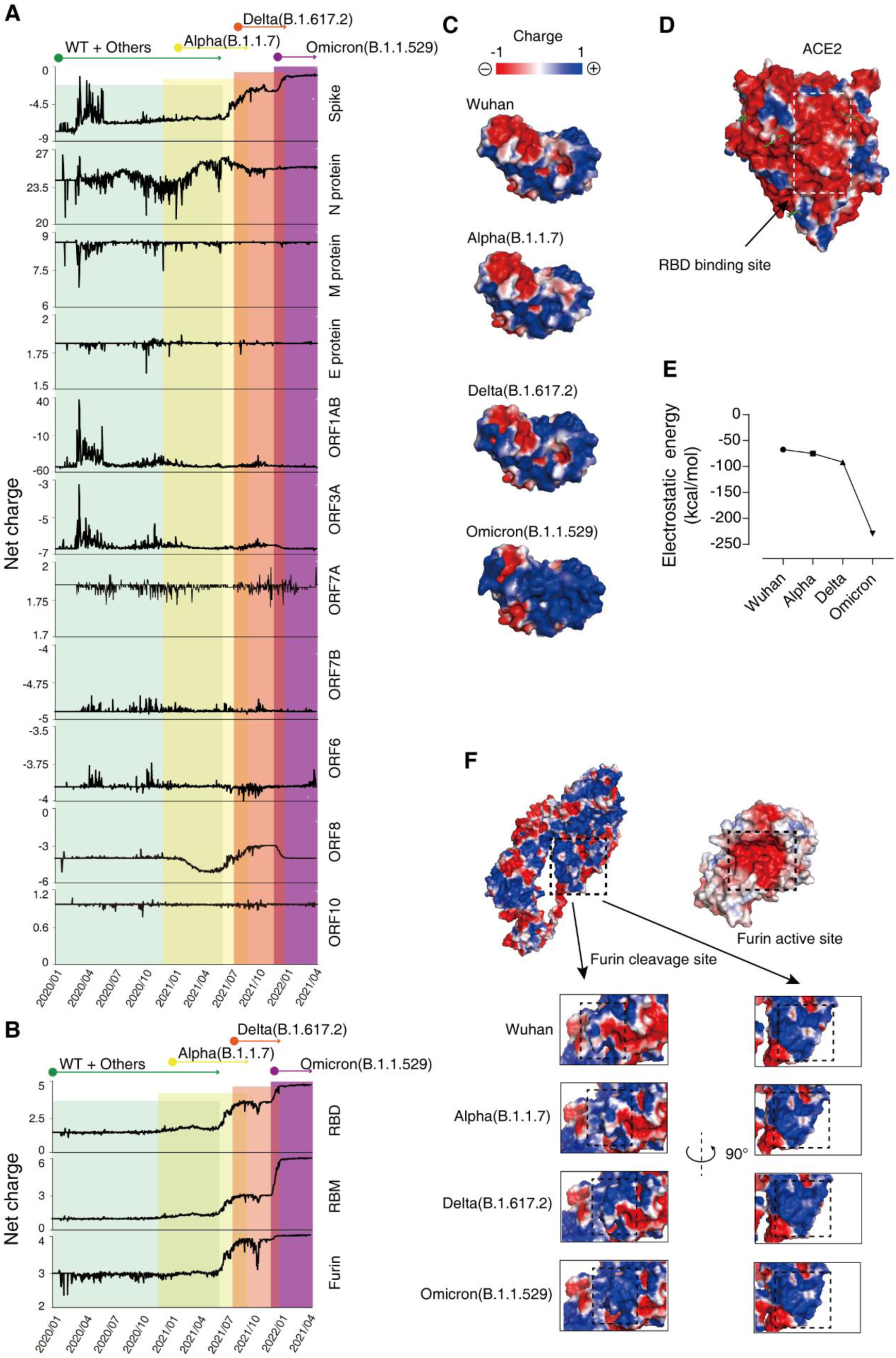
The charge of SARS-COV-2 proteins during evolution. (A) Net charge evolution of SARS-COV2 proteins. (B) Net charge of local region of the spike protein: RBD (333-527), RBM (438-506) and Furin cleavage site (675-695) [27,28]. (C, D) Electrostatic surfaces of the spike protein RBD region and human ACE2 protein. (E) Electrostatic energy of ACE2 binding S-RBD. (F) Electrostatic surfaces of the spike protein furin cleavage site and human furin.

Indeed, the human ACE2 protein is largely negatively charged in surface, especially in the region that binds SARS-COV-2 S-RBD domain (Fig. 3C). The increasing positive charge in RBD (Fig. 3D) obviously elevated the electrostatic affinity, confirmed by energy computation on the structure of the RBD-ACE2 binding state (Fig. 3E). Similarly, the furin enzyme active site is highly negatively charged. The furin cleavage site in S-gene accumulated positive charges during evolution (Fig. 3F), which should increase the binding efficiency and accelerate the enzymatic cleavage.

## 4. DISCUSSION

Our results may provide novel insights on the SARS-COV-2 evolution. It has been shown that the narrow-spectrum +ssRNA viruses evolve their codon usage matching their hosts’ tRNA better than the broad-spectrum viruses. Such adaptation is to optimize their protein expression [29,30]. In our analyses, we showed that the SARS-COV-2 did not evolve towards optimized codon usage in human lung cells, indicating that it maintains the ability to infect multiple hosts. It has been experimentally shown that the SARS-COV-2 virus infects cats and dogs [31,32] Such ability facilitates formation of a natural reservoir of virus. Even with the strictest lockdown policy, the reservoir serves as a refuge of SARS-COV-2 and may transmit to human again when possible. Indeed, the SARS-COV-2 virus is believed to transmit initially from infected animal to human in a wild animal market in Wuhan, China [33]. Recently, after about 3 weeks of strict lockdown policies in Shanghai, the number of new confirmed cases did not drop, suggesting that the virulence of such a natural reservoir of the omicron variant could not be neglected. Maintaining a broad spectrum of hosts facilitates the survival of SARS-COV-2, especially in the regions deploying “dynamic zero-COVID policy” like China. In other countries where not so strict policies were applied, the massive vaccination also drives the virus to maintain the ability to infect other species to survive.

Therefore, it is understandable that the major evolutionary driving force is applied on the infectivity, i.e. the spike protein. Coincidently, we showed that the S gene is the only gene which underwent the positive selection. Since the infection is conducted via the binding of S-RBD and human ACE2, two major ways can enhance this affinity: increasing the electrostatic interaction and mutating specific residues to form more stable van der Waals force on the interaction interface. Electrostatic force decreases far slower than the van der Waals force against distance, making it more suitable to serve as guide force of the two molecules. Increasing the electrostatic interaction is also universal for all species, facilitating the maintenance of the broad host spectrum. In contrast, mutating specific residues to form more stable van der Waals force is highly host-specific, because the structure of the ACE2 in different species vary remarkably. Therefore, increasing the electrostatic interaction becomes the best evolutionary strategy of the SARS-COV-2 virus for higher infectivity.

With the average reported R_0_=9.5 [34], the SARS-COV-2 omicron variant is already among the most infectious viruses in the human history. However, since the accumulation of positive charges in its S-RBD and Furin digestion site is the key aspect of the high infectivity, it can be used as druggable target. Polar reagents which neutralize the charges may counteract the infectivity. In another aspect, the positive charges cover almost the entire interaction surface of the S-RBD and Furin digestion site of the omicron variant, showing that there’s only restricted potential to accumulate more positive charges in these interfaces, which indicates that the infectivity of SARS-COV-2 may approach to a limit.

## Author Contributions

Xiaolong Lu: Methodology, Software, Data curation and analysis, Visualization, Writing. Gong Zhang: Conceptualization, Project administration, Funding acquisition, Supervision, Visualization, Data curation, Writing - original draft, Writing - review & editing.

## Data Availability Statement

The data presented in this study are available on accession number

